# Methylome dynamic upon proteasome inhibition by the *Pseudomonas syringae* virulence factor Syringolin A

**DOI:** 10.1101/2021.01.19.427327

**Authors:** DMV Bonnet, S Grob, L Tirot, PE Jullien

## Abstract

DNA methylation is an important epigenetic mark required for proper gene expression and silencing of transposable elements. DNA methylation patterns can be modified by environmental factors such as pathogen infection, where modification of DNA methylation can be associated with plant resistance. To counter the plant defense pathways, pathogens produce effectors molecule, several of which act as proteasome inhibitors. Here we investigated the effect of proteasome inhibition by the bacterial virulence factor Syringolin A on genome-wide DNA methylation. We show that Syringolin A treatment results in an increase of DNA methylation at centromeric and pericentromeric regions of *Arabidopsis* chromosomes. We identify several CHH DMRs that are enriched in the proximity of transcriptional start sites. Syringolin A treatment does not result in significant changes in small RNA composition. However, significant changes in genome transcriptional activity can be observed, including a strong upregulation of resistance genes that are located on chromosomal arms. We hypothesize that DNA methylation changes could be linked to the upregulation of some atypical members of the *de novo* DNA methylation pathway: *AGO3*, *AGO9* and *DRM1*. Our data suggests that modification of genome-wide DNA methylation resulting from an inhibition of the proteasome by bacterial effectors could be part of a epi-genomic arms race against pathogens.

---

Plants are sessile organism and must constantly adapt their gene expression to react to constantly changing environments. Among other environmental stresses, the infection of plants by pathogens, such as bacteria or fungi is a constant threat and leads to significant losses in crop production. Plants have evolved several ways to defend themselves against pathogens, starting from physical barriers, such as the cuticula, to specific intra-cellular pathways that recognize pathogen effector molecules. The effector-triggered immunity pathway (ETI) involves the specific recognition of effector molecules that are secreted by the pathogen and counteract the plant primary immune response (Jones and Dang, 2006).

Interestingly, several effector molecules produced by bacteria, viruses, and fungi were found to inhibit the host proteasome machinery (Chiu et al., 2010; Dudler, 2013; Groll et al., 2008; Jin et al., 2007; Marino et al., 2012; Sahana et al., 2012; Sorel et al., 2019; Üstün and Börnke, 2014; Üstün et al., 2016; Verchot, 2016) and - more generally - to regulate protein degradation (Langin et al., 2020). Thus, proteasome inhibition seems to represent a common strategy used by plant pathogens. Additionally, host proteasome inhibition has been shown to suppress systemic acquired resistance (SAR) and to consequently increase plant susceptibility to bacterial pathogen (Üstün et al., 2016). Beyond classical bacterial type III effector such as XopJ and HopM1 (Üstün et al., 2013; Üstün et al., 2016), which inhibit proteasome activity by interacting with proteasome subunits, a virulence factor called Syringolin A (SylA) inhibits the proteasome via an irreversible covalent binding to the proteasome catalytic subunits (Groll et al., 2008). SylA is a small tri-peptide derivative, which is secreted by the bacteria *Pseudomonas Syringae pv syringae* (*Pss*). Bacteria deficient for SylA secretion are less virulent than their wild-type counterpart. SylA secretion is important for the bacteria to overcome stomata closure and consequently gaining better leaf penetration (Schellenberg et al., 2010). Similarly, SylA was also found to be important for wound entry and colonization of the host via the vascular tissue (Misas-Villamil et al., 2013). Interestingly, SylA was found to accumulate in the nucleus, which lead the authors to suggest it could act preferentially on the nuclear proteasome (Kolodziejek et al., 2011).

The proteasome is one of the main protein degradation machinery in eukaryotic cells. Protein targeting to the proteasome relies in part to poly-ubiquitination by E3 ubiquitin ligases (Sadanandom et al., 2012). Due to its central role in the cell, the proteasome machinery is involved in several biological processes, such as hormonal signaling, circadian clock as well as plant stress response (Sadanandom et al., 2012; Vierstra, 2009). Additionally, the proteasome has also been linked to the regulation of chromatin and transcription (Geng et al., 2012). The proteasome’s role in the regulation of chromatin has not only been linked to the regulation of chromatin components, such as histones and their assembly into nucleosomes, but also chromatin regulators, such as histone chaperones and histone modifiers. (Bach and Hegde, 2016; H. Karimi Kinyamu et al., 2008; Jeong et al., 2011; Lee et al., 2011). In addition, proteins involved in DNA methylation were found to be regulated by the proteasome.

DNA methylation in *Arabidopsis* is characterized by the apposition of a methyl group to cytosine residues. In plants, DNA methylation occurs on three distinct sequence contexts: CG, CHG and CHH, where H stands for any nucleotide except cytosine. Each methylation context is regulated by its own cognate pathway (Law and Jacobsen, 2010). CG methylation is maintained by the maintenance DNA methyltransferase MET1 (METHYLTRANSFERASE 1). Despite the lack of evidence for a proteasome regulation of the MET1 protein, other important proteins regulating CG methylation, such as the VIM proteins are E3 ubiquitin ligases (Johnson et al., 2007; Kraft et al., 2008; Woo et al., 2007), suggesting a link between ubiquitination and CG methylation. The main enzyme linked to CHG methylation is the DNA methyltransferase CMT3 (CHROMOMETHYLASE 3) (Lindroth and Jacobsen, 2001). CMT3 itself was found to be targeted for proteasomal degradation by an atypical JmjC domain protein, JMJ24, which has E3 ubiquitin ligase activity (Deng et al., 2016). Recently, DRM2 (DOMAINS REARRANGED METHYLASE 2), the main DNA methyltransferase involved in *de novo* cytosine methylation at CHH sites was also found to be regulated by an E3 ubiquitin ligase named CFK1, leading to its targeting for degradation by the proteasome (Chen et al., 2020). DNA methylation in all sequence contexts can, therefore, be directly or indirectly linked to ubiquitin-dependent proteasome degradation.

DNA methylation has previously been shown to be involved in the response against bacterial and fungal pathogens (Deleris et al., 2016; Zhu et al., 2016). Indeed, mutants with lower DNA methylation levels, such as *met1* were shown to display an increased resistance to the bacteria *Pseudomonas* (Dowen et al., 2012), while mutant displaying global DNA hypermethylation such as *ros1* were shown to be more susceptible to *Pseudomonas*, *Fusarium,* and *Hyaloperonospora* infection (Le et al., 2014; López Sánchez et al., 2016; Schumann et al., 2019; Yu et al., 2013). DNA *de novo* methylation is known to be targeted by small RNA molecules of principally 24 nucleotides (nt) in size. This pathway is referred as RNA-directed DNA methylation (RdDM) (Matzke and Mosher, 2014). Several members of the RdDM pathway were also found to be involved in *Botrytis*, *Plectosphaerella,* and *Pseudomonas* resistance (Agorio and Vera, 2007; López et al., 2011).*Pseudomonas* bacterial infection was shown to affect genome-wide DNA methylation pattern (Dowen et al., 2012). Additionally, DNA methylation is involved in transgenerational memory of bacterial infection, a phenomenon referred as Transgenerational Acquired Resistance (TAR) (Luna and Ton, 2012; Luna et al., 2012).

Noting the converging evidence for a role of DNA methylation during bacterial infection as well as proteasome inhibition during this process, we sought to investigate the effect of proteasome inhibition on global DNA methylation levels. Here we show that genome-wide DNA methylation is affected by treatment with the proteasome inhibitor SylA. SylA treatment results in a moderate genome hypermethylation in all sequence context. This hypermethylation is mainly affecting centromeric region of *Arabidopsis* chromosomes. We identified 10101 100bp DMRs of which 6304 are CHH-hypermethylated DMRs. We could not identify major changes in small RNA composition upon SylA treatment suggesting that small RNAs might not have a direct effect on the observed DNA methylation changes. Furthermore, the *Arabidopsis* transcriptome is strongly affected by SylA-mediated proteasome inhibition. We propose that the induction of atypical members of the RdDM pathway such as *DRM1*, *AGO9* and *AGO3* might explain the changes observed upon SylA treatment.

## RESULTS

### Syringolin A induces moderate centromeric hypermethylation

In order to investigate the effect of proteasome inhibition on the methylome of *Arabidopsis*, we performed genome-wide bisulfite sequencing (GWBS) of *Arabidopsis* plantlets treated with the bacterial virulence factor SylA, a known proteasome inhibitor (Groll et al., 2008). Considering all three sequence contexts, we obtained DNA methylation levels for 14269258 and 10867987 cytosines for mock- and sylA-treated GWBS libraries, respectively, that reached more than 10-fold coverage. The obtained data were plotted as a circular heat map using 50 kb genomic bins (Fig 1A). We observed that the distribution of the genome-wide DNA methylation did not change between treated and the control samples in all sequence context CG, CHG and CHH (Fig 1A). As previously observed (Cokus et al., 2008; Lister et al., 2008; Stroud et al., 2012), DNA methylation is higher in centromeric and pericentromeric regions of the *Arabidopsis* genome, coinciding with increased transposon densities. As expected, the percentage of DNA methylation is higher in CG context (28.36% and 29.25%), then CHG context (12.05% and 13.62%) and finally at its lowest in CHH context (3.16% and 3.54%) in both control and treated samples. In order to refine our analyses, we focused on methylated windows (i.e., >40% methylation for CG, >20% for CHG and >10% for CHH) (Fig 1B). We could observe a slight but consistent increase in DNA methylation levels upon SylA treatment in all sequence contexts. In order to investigate in which chromosomal environment this increase of DNA methylation occurs, we plotted the percent of changes in DNA methylation across the genome. We observed that, except for a few discreet loci on chromosome arms, most of the increase in DNA methylation occurred in the centromeric and pericentromeric regions of all *Arabidopsis* chromosomes (Fig 1C and Fig S1). We conclude that proteasome inhibition by SylA treatment does not result in a global reprograming of genome-wide DNA methylation but rather in a moderate hypermethylation of the centromeric region.

**Fig.1.**
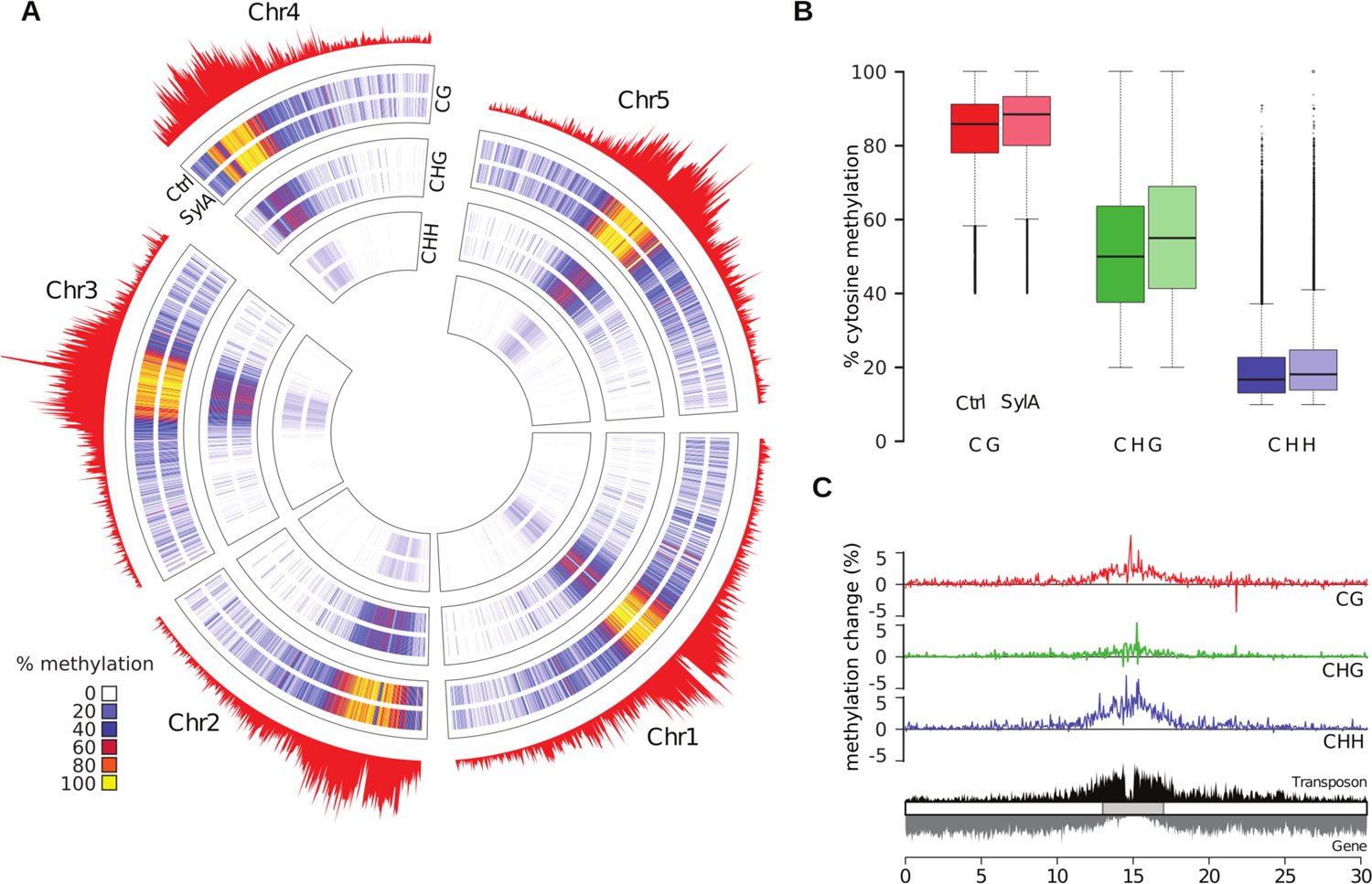
Methylation changes upon Syringolin A treatment. A) Circos plot for cytosine methylation levels in CG, CHG, and CHH contexts in mock treated (Ctrl) and Syringolin A treated (SylA) plantlets. Red barplots depict transposable element densities within 50 kb genomic bins. B) Boxplot showing global cytosine methylation levels using 100 bp bins. Methylated bins were defined by a minimal methylation level of 40 % for CG context (Red), 20 % for CHG context (Green), and 10 % for CHH context (Blue), respectively. Lighter colour represent the SylA treated sample. C) Percent methylation changes along chromosome 1 for CG (red), CHG (green), and CHH (blue) context, respectively (See also Fig.S1). Bottom track depicts transposable element (black) and gene (grey) density in 50 kb bins as a proxy for the occurrence of heterochromatin and euchromatin. Grey rectangle: pericentromere.

### Syringolin A treatment results mainly in CHH DMRs

To investigate the changes in DNA methylation on local scale, we identified differentially methylated regions (DMRs) of 100 bp in size for each DNA methylation context. To satisfy the criteria as a DMR, the respective 100 bp genomic bins had to differ by at least 10 % between treatments and exhibiting a q-value < 0.05. We identified a total of 10101 individual genomic bins satisfying DMR criteria (Fig 2A). Most DMRs were in CHH context (9173) compared to CG (409) and CHG (672). Additionally, they were mostly hyper DMRs (gain of DNA methylation) rather than hypo DMRs (lost of DNA methylation). CHH DMRs were mostly corresponding to transposable elements (TEs) (51%), UTRs (14%), promotors (14%), and intergenic sequences (15%) and were rather depleted in genic sequences (2%) (Fig 2B). Similarly, CG and CHG DMRs could also majorly be associated to TEs (Fig S2). The overlap between the DMRs of different contexts is limited but the overlap is significant (permutation-based *p*-value using 10^4^ repetitions for all dual overlaps and triple overlaps < 10^-4^). Taking into account the frequent association of DMRs of all contexts with TEs, we observed an especially pronounced overlap when regarding DMR - associated TEs (Fig S3), suggesting that most significant changes in DNA methylation occurs within or at proximity of TEs. In order to visualize in which genomic region DMRs are located, we plotted the DMRs along the five *Arabidopsis* chromosomes (Fig 2C). Both hypo and hyper CHH DMRs are enriched in centromeric and pericentromeric region mirroring the TE density distribution in the *Arabidopsis* genome. In order to better characterize the DMRs location and their potential effect on gene expression, we computed the distribution of their mean distance to transcriptional start sites (TSS) and compared it to that of non-DMR methylated regions. For this purpose, we separated our analysis between the DMRs occurring in the pericentromeric regions and the DMRs occurring on chromosomal arms. Interestingly, SylA induced DMRs in chromosomal arms are closer to TSS compared to randomly sampled methylated regions for both DMRs associated with TEs (Fig 2D) and genes (Fig 2E). This increased proximity to TSS is more pronounced for CHH DMRs (Fig 2D-E) than for CG and CHG DMRs (Fig S4A-B). No significant differences were observed for DMRs situated in pericentromeric regions (Fig S4 C-D). In summary, our results show that SylA treatment results principally in CHH DMRs situated in centromeric and pericentromeric region and that DMRs situated in chromosomal arms are in close proximity to TSS.

**Fig.2.**
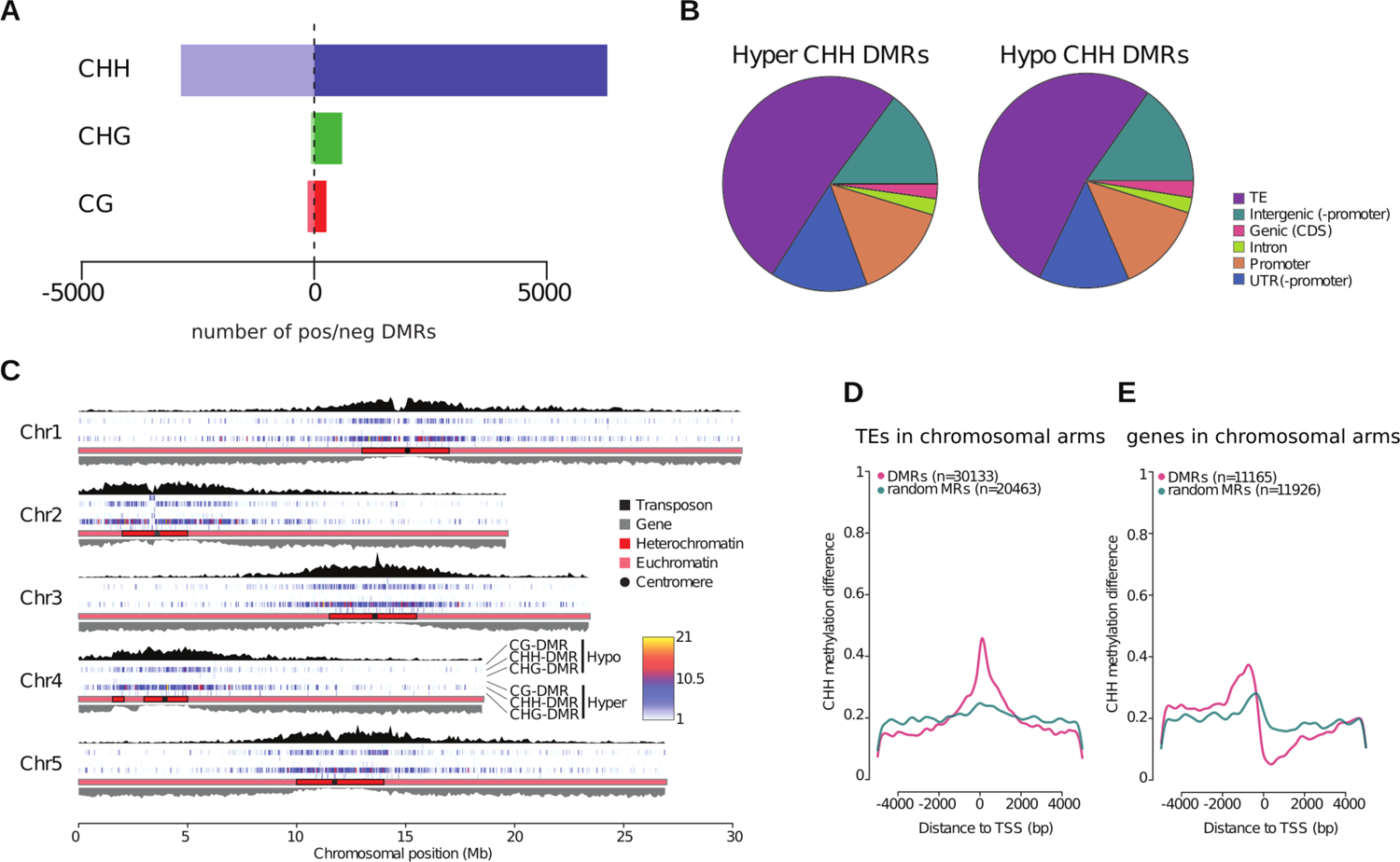
DMRs induced by SylA treatment. A) Number of hyper and hypo DMRs per context. B) Pie chart representing genomic features, associated with CHH DMRs (taking 1 kb 5’ of the annotation start and 3’ of the end position, respectively). Left: CHH hyper DMRs, right: CHH hypo DMRs. Genomic features not visible on the pie chart due to low occurrence of DMRs have been omitted. C) Genomic localization of DMRs. The colour code indicates the number of DMRs per 50 kb genomic bin. Transposable element (TE) (black) and gene density per 100 kb bin are depicted as a proxy for the occurrence of heterochromatin and euchromatin, respectively. D) Metaplot showing the mean distance from CHH DMRs to the putative transcriptional start site (TSS) of TEs located on chromosome arms in closest proximity to the DMR. DMRs within 5 kb of TEs’ annotation start sites were selected. Magenta: DMRs, green: randomly selected non-differentially methylated regions occurring within 5 kb of TEs’ annotation start sites. E) Metaplot showing the mean distance from CHH DMRs to the TSS of genes located on chromosome arms in closest proximity to the DMR. DMRs within 5 kb of genes located on chromosome arms were selected. Magenta: DMRs, green: randomly selected non-differentially methylated regions occurring within 5 kb of TSSs.

As previously mentioned, DNA methylation changes were observed upon *Arabidopsis* infection by the bacteria *Pst* (Dowen et al., 2012). In order to evaluate how similar SylA DMRs are from *Pst* induced DMRs, we analyzed the overlap between *Pst* and SylA DMR associated features (Fig S8A-C)). Although no significant overlap could be observed for CG DMRs or CHG DMRs (Fig S8A-B), a significant overlap could be observed concerning CHH DMRs with 413 overlapping loci (*p*-value < 10^-4^) (Fig S8C). The overlap between *Pst* and SylA DMRs suggests that these DMRs could be linked to proteasome inhibition taking place during *Pst* infection. Further studies will be required to confirm this hypothesis.

### Syringolin A treatment does not influence small RNA population

DNA methylation in CHH context is closely link to the RNA-directed DNA methylation pathway (RdDM). This pathway relies on small RNAs to induce DNA methylation at the complementary DNA locus. In its canonical form, RdDM relies on the loading of 24nt small RNAs by AGO4 or AGO6 and the recruitment of the *de novo* methyltransferase DRM2 to the DNA (Law and Jacobsen, 2010; Matzke and Mosher, 2014). In order to investigate potential changes in small RNA populations between mock-treated and SylA-treated plantlets, we profiled the small RNA populations using next-generation sequencing. As expected in both conditions, 24nt small RNAs comprise the most abundant size fraction and are mostly starting with a 5’A and are predominantly mapping to TEs (Fig3A-C). In both samples, the second most abundant are 21nt small RNAs. They are mainly starting with a 5’T and likely correspond to microRNAs. 22nt small RNA are the least abundant of the three sizes, predominantly starting with a 5’G and mapping to TEs. We conclude that the overall small RNA composition is very similar between the control and the treated samples and comparable to previously published *Arabidopsis* small RNA compositions (Mi et al., 2008). To investigate the small RNA population in more detail, we selected 100bp genomic bins showing differential expression (exhibiting at least 10 reads across the two treatments and showing at least a 4-fold change between the treatments). We found that despite being very similar, several bins (n=1539) mostly comprising 24nt small RNAs (n=1016) were enriched in the SylA-treated sample (Fig 3D). Bins enriched in 21nt and 22nt small RNA were not as abundant (21 nt: 326, 22 nt: 197) (Fig 3D). For all sRNA sizes we identified only a low number of sRNA depleted bins (n=192). As expected, the enriched 24nt sRNAs were predominantly starting with a 5’A and mapping to TEs (Fig S9A-C). The enriched 21nt and 22nt small RNA had an unusual 5’C and 5’T enrichment, respectively, and were predominantly mapping to TEs (Fig S9A-C). In comparison to the total library populations, 21 nt sRNA, presumably representing microRNAs, were underrepresented in the SylA enriched sRNA population, indicating that micro RNAs are not activated upon SylA exposition. Bins enriched in 24nt sRNA are located principally to pericentromeric regions (Fig S5).

**Fig.3.**
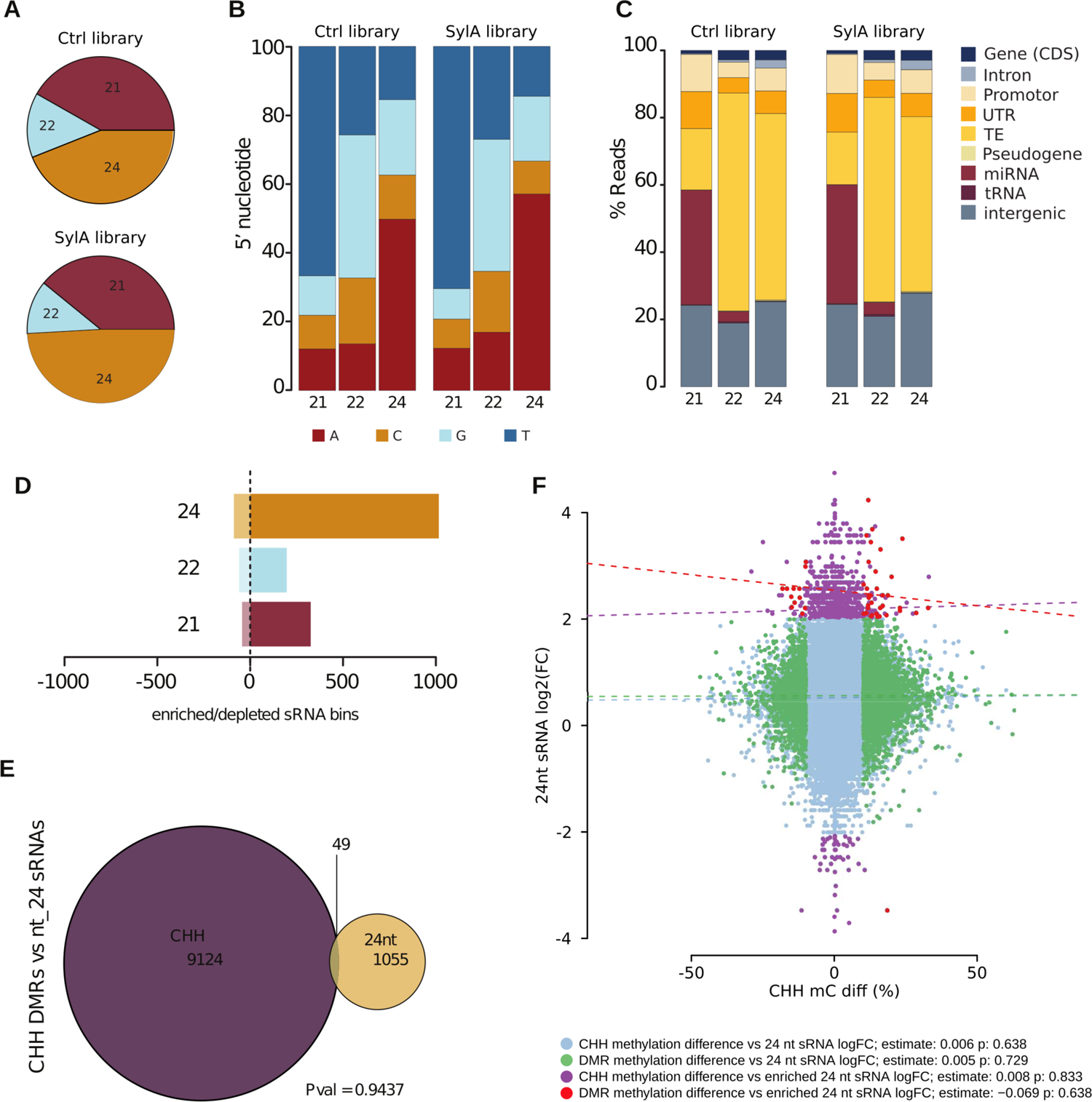
small RNA population upon Syringolin A treatment. A) Pie charts representing sRNA size distribution (21 nt, 22 nt, 24 nt) in total mock (Ctrl) treated libraries and Syringolin A (SylA) treated libraries. B) Stacked bar charts showing the percentage of the 5’ nucleotide identity by sRNA sizes. C) Genomic features associated with sRNAs per size (21 nt, 22 nt, 24 nt). small RNA bins lying within 1 kb 5’ of the annotation start site and 1 kb 3’ of the annotation end site were selected. D) Barplot featuring the number of enriched and depleted 100 bp small RNA bins upon SylA treatment. E) Venn diagram showing the overlap between CHH 100 bp DMRs and enriched/depleted 24 nt 100 bp sRNA bins. F) correlation analysis between methylation levels of DMRs with 24nt enriched Bins showing a lack of correlation.

However, only 49 of the 1055 24nt enriched bins overlapped with CHH DMR bins (Fig 3E, p=0.9437). The same observation was made for 21nt and 22nt enriched small RNA bins (Fig S6D-F). Additionally, we could not find a significant correlation between the increase in 24nt sRNAs and the CHH methylation levels (Fig 3F). The absence of significant correlation was also observed for 21nt and 22nt enriched bins (Fig S6A-C). We conclude that the small RNA composition is mostly unaffected by the SylA treatment and that the moderate changes in sRNA composition or abundance do not correlate with the changes in CHH DNA methylation and is likely not the cause of this changes.

### Syringolin A treatment predominantly affects transcription in chromosomal arms

DNA methylation is a well-known regulator of gene and transposon expression. In order to investigate a potential link between the observed changes in DNA methylation upon SylA treatment and genome-wide transcriptional activity, we subjected our samples to RNA sequencing. Upon SylA treatment, we could detect a major change in transcriptional activity (Fig 4A-B), identifying a total of 3241 differentially expressed transcripts (adjP < 0.01 and abs(logFC) > 2), among them 2945 differentially expressed genes (DEG). We noted a bias toward upregulated DEGs (1675) compared to downregulated DEGs (1270), showing that SylA treatment rather leads to transcriptional up-regulation than down-regulation. Interestingly, gene ontology (GO) term enrichment analysis for the upregulated DEGs revealed an enrichment in “positive regulation of RNA polymerase II initiation” (GO:0045899, pVal= 9.4*10^-6^) (Fig 4D, Table S1). Our results suggest that an inhibition of the proteasome using SylA results in an increased transcriptional activity. We can observe a similar GO term enrichment in “positive regulation of RNA polymerase II initiation” (GO:0045899, pVal= 2.7*10^-9^) using previously published transcriptomic data using MG132, a non-covalent proteasome inhibitor (Table S1) (Gladman et al., 2016). This suggests that this feature in not only linked to SylA itself but more generally to proteasome inhibition. Increased transcriptional activity in response to proteasome inhibition seems to be widely conserved, as it was also observed in human cells treated with MG132 (Kinyamu et al 2019).

**Fig4.**
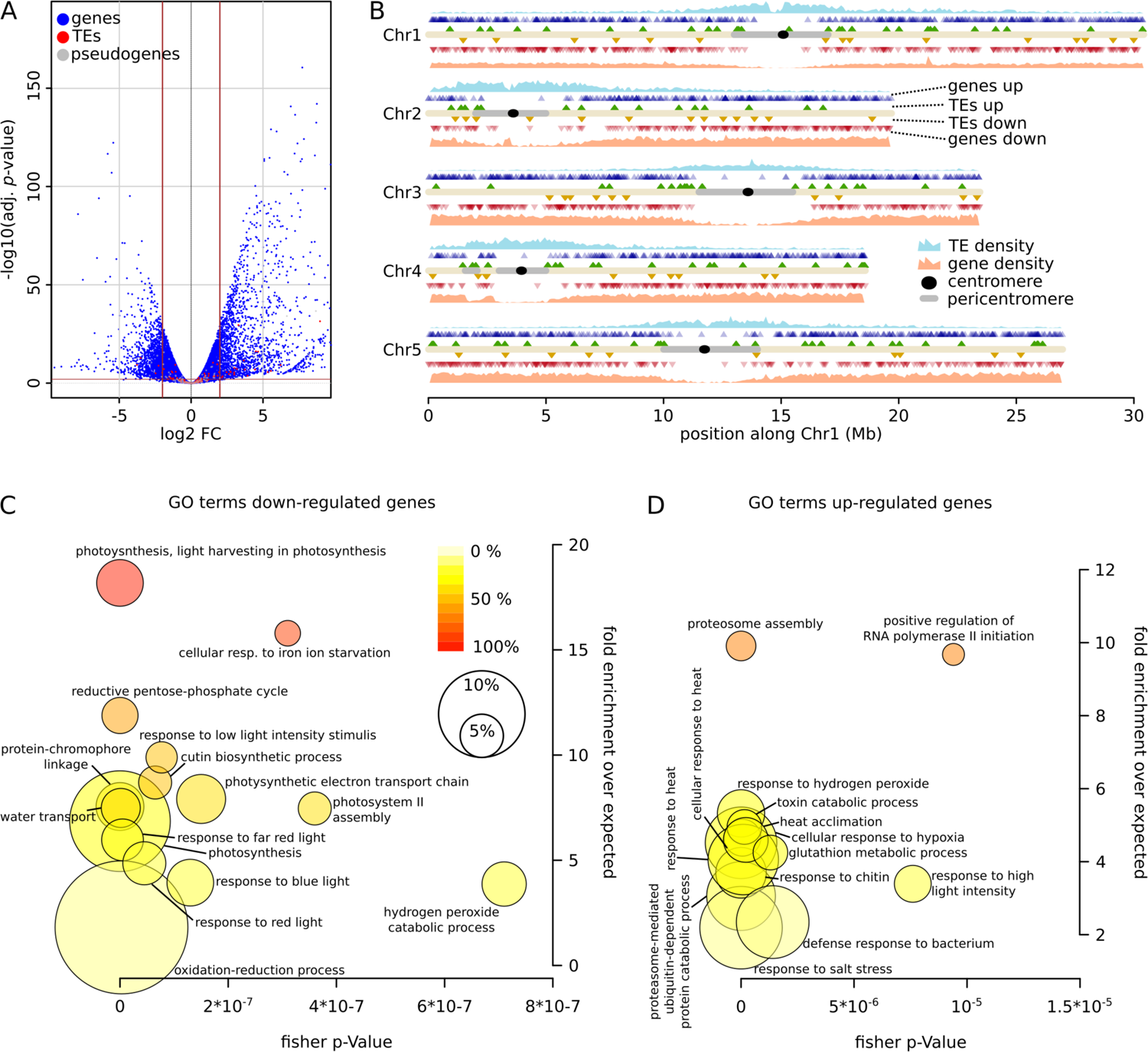
High number of up-regulated genes upon Syringolin A treatment. A) Volcano plot depicting up- and down-regulated genes (blue), TEs (red), and pseudogenes (grey). Brown horizontal and vertical lines mark the thresholds to call significantly differential expression (logFC > 2, adjusted *P*-value < 0.01). B) Chromosomal location of significantly up-regulated genes (blue triangles) and TEs (green triangles) and down-regulated genes (red triangles) and TEs (orange triangles). Gene and TE density in 50 kb bins are shown as a proxy for the occurrence of euchromatin and heterochromatin. (C-D) GO term analysis for down-regulated (C) and up-regulated genes (D). Most significant 15 GO terms are shown. Bubble size indicates the percentage of all down-regulated genes within the respective GO term. Colour code illustrates the percentage of significant genes of all genes of the given term. Complete GO Term enrichment list can be found in Table S1.

As previously reported upon SylA treatment (Michel et al., 2006), we observed an enrichment in “proteasome assembly” (GO:0043248, pVal= 10^-9^). We hypothesize that in order to compensate for the inhibition of the proteasome, proteasome related genes are transcribed at higher levels. This phenomena was also observed using MG132 (Table S1) (Gladman et al., 2016). Similarly, an increase of proteasome-related gene expression also occurs during *Pst* infection (Üstün et al., 2018), further supporting that proteasome inhibition genuinely happen during *Pst* infection. In addition, up-regulated DEGs are enriched in GO terms describing several abiotic and biotic stresses (Fig 4D). Down-regulated DEGs are clearly enriched in GO terms related to photosynthesis as well as response to light (Fig 4C, Table S1). A link between ubiquitin-dependent proteasome degradation and light response in Arabidopsis has been long established with the discovery in plant of the COP9 signalosome (Chamovitz et al., 1996; Cope and Deshaies, 2003; Wei and Deng, 2003).In addition to modifying gene expression, SylA treatment also affects the expression of TEs, 217 of them exhibiting significantly differential expression (Fig 4A-B), with a strong tendency towards being upregulated rather than downregulated (up: 145, down:72). Surprisingly and similarly to DEGs, we predominantly detected differentially expressed TEs located on chromosomal arms rather than pericentromeric or centromeric region (Fig 4B), despite the majority of TEs being located there. Upregulated TEs are enriched in LTR-copia TE families (Fig S7), whereas downregulated TE tend to be enriched in DNA-MuDR transposons (Fig S7). Our result shows that SylA treatment has an important effect on *Arabidopsis* transcriptome, which is likely linked to role of SylA as a proteasome inhibitor.

### Syringolin A treatment induces the expression of atypical DNA methylation factors

In order to understand how SylA could influence DNA methylation beyond its role in inhibiting the proteasome, we investigated if it could act by modulating the expression of key actors of DNA methylation. We have focused our analyses on the key members of the different DNA methylation pathways and homologous genes encoded in the *Arabidopsis* genome (Fig 5A). Methylation on CG sites requires the principal CG DNA methyltransferase *MET1* (Ennis et al., 1996; Kankel et al., 2003; Saze et al., 2003). Additionally, on centromere and pericentromeric sequences, the chromatin remodelers *DDM1* and *MOM1* are required for the maintenance of CG methylation (Jeddeloh et al., 1998; Zemach et al., 2013). In addition, the *Arabidopsis* genome encodes 3 other *MET* genes: *MET2a*, *MET2b* and *MET3* (Jullien et al., 2012). None of these proteins affecting CG methylation showed a sufficient expression fold-change to meet DEG criteria. Similarly, the CHG DNA methyltransferases (*CMT1*, *CMT2* and *CMT3*) (Henikoff and Comai, 1998; Lindroth et al., 2001) and the DNA demethylase (*ROS1*, *DME*, *DML2* and *DML3*) (Choi et al., 2002; Gong et al., 2002; Penterman et al., 2007) were not among DEGs. However, we observed a significant upregulation for some members of the *de novo* DNA methylation RdDM pathway, which elicits CHH methylation. We analyzed the *de novo* DNA methyltransferases *DRM1*, *DRM2* and *DRM3*. We could observe a significant induction of *DRM1* (log2(FC)=5.9). A significant increase in transcript abundance could be observed from 8h post induction. In the RdDM pathway, small RNA molecules, loaded in protein called Argonautes, are targeting *de novo* methylation on the complementary DNA sequence. The Argonautes associated we the RdDM pathway are *AGO4*, *AGO6*, *AGO9*, *AGO8* and *AGO3* (Duan et al., 2015; Havecker et al., 2010; Huang et al., 2016; Stroud et al., 2012; Zilberman et al., 2003). We found that *AGO3* (log2(FC)=3.4) and *AGO9* (log2(FC)=2.2) were significantly upregulated in response to SylA treatment. Performing quantitative PCR on *Arabidospis* cDNA, we could confirm that SylA treatment leads to an upregulation of *AGO3*, *AGO9* and *DRM1* (Fig 5B).

**Fig5.**
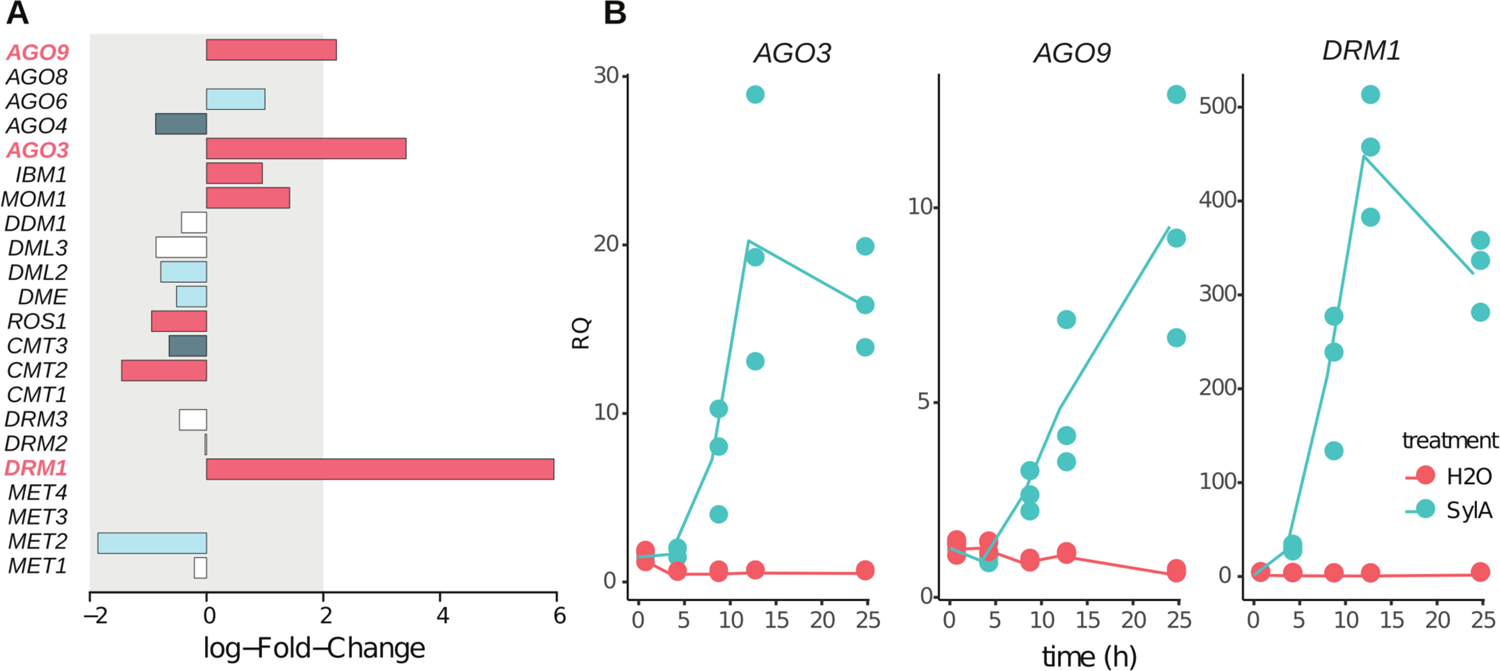
**Syringolin A regulate the expression of AGO3, AGO9 and DRM1**. A) Histogram showing the log Fold change of *Arabidopsis* genes involved in the DNA methylation pathway. The bars are colored according to the FDR value (Red: FDR<0.0001, dark blue: FDR[0.001-0.0001], light blue: FDR[0.01-0.001] and white: FDR>0.01). B) Quantitative PCR showing the upregulation of *AGO3*, *AGO9* and *DRM1* during a time course following a treatment of seedlings with SylA or H2O as control. Individual points represent biological replicates, line represent the Mean RQ value. ACT2 was used as normalizer.

## DISCUSSION

Our study shows that Syringolin A, a virulence factor secreted by *Pseudomonas Pss* strains, induces moderated change of the *Arabidopsis* methylome. Differentially methylated regions are mostly located in the centromeric and pericentromeric regions of the *Arabidopsis* chromosomes. Upon SylA treatment, substantial changes in the *Arabidopsis* transcriptome are observed, which are most likely linked to SylA action on the proteasome rather than on its effect on DNA methylation. We identified potential candidate genes that could be involved in the observed increase in DNA methylation levels. These candidates are atypical members of the RNA directed DNA methylation pathway: *AGO3*, *AGO9* and *DRM1*.

Beyond SylA treatment, dynamic changes in centromeric DNA methylation seems to also occur during *Arabidopsis* infection by *Pseudomonas synringae pv. tomato* (*Pst*), which in contrast to *Pseudomonas synringae pv synringae* (*Pss*) does not secrete SylA. Changes in DNA methylation levels during *Pseudomonas* infection were previously reported (Dowen et al., 2012; Pavet et al., 2006). Pavet *et al* showed that DNA methylation decreases in centromeric region during early infection (1 day post infection) with *Pst* (Pavet et al., 2006), whereas Dowen *et al* report an increase of centromeric DNA methylation at five days post infection. At 5 days post infection, similar to SylA treated plantlets, the increase of DNA methylation was clear in the CHG and less pronounced in CG and CHH context. Despite not secreting SylA, *Pst* also uses type III effectors, such as HopM1, to inhibit the proteasome during infection (Üstün et al., 2016). It is therefore likely that at least parts of the DNA methylation changes observed during *Pst* infection, at 5 days after infection, could be linked to proteasome inhibition by type III effectors as we have observed using SylA. Considering the temporal effect of the infection on centromeric DNA methylation, one may speculate that an early response of plants to infection triggers DNA demethylation, as suggested by centromeric demethylation at 1dpi and the preferential reactivation and, thus, upregulation of transposable elements (TEs) observed upon Flagellin treatment (Yu et al., 2013). This DNA demethylation has been associated with the expression of NBR/LRR genes which are often in close proximity to TEs. At later stages of infection, however, the action of bacterial effectors, such as proteasome inhibitors, might counter act this DNA demethylation and induce centromeric hypermethylation.

Interestingly, in contrast to the changes in DNA methylation mainly occurring in centromeric and pericentromeric regions, we observed the majority of differentially expressed TEs and DEGs to be located on chromosomal arms. If linked, such phenomena would imply a *trans* effect of the centromere DNA methylation and/or organization on the expression of genes situated on the chromosomal arms. Such a potential *trans* effect has been noted previously in response to *Pst* infection (Cambiagno et al., 2018). Indeed, loss of centromeric and pericentromeric DNA methylation in early infection has been hypothesized to be linked to the activation of NBS/LRR disease resistance genes situated on chromosomal arms (Cambiagno et al., 2018). It was hypothesized that centromeric hypomethylation would lead to a recruitment of the RdDM RNA machinery to the centromere in order to remethylate centromeric TE dense regions. As a consequence, the RdDM machinery would be depleted from the chromosomal arm and would therefore facilitate PRR/NLR genes expression. Another hint for a *trans* effect during infection came from epigenetic quantitative trait loci (QTL) linked with increased resistance to the fungi *Hyaloperonospora.* These epigenetic QTLs are located in pericentromeric regions but are associated to increased priming of resistance genes situated on chromosomal arms (Furci et al., 2019). It was hypothesized that this *trans*-effect might be due to changes in the 3D organization of the genome that would affect genes transcriptional activity. However, to date, it is still not fully clear how centromeric and pericentromeric DNA methylation and/or organization would affect the expression of resistance genes *in trans*. Further studies will be required to evaluate if this *trans* effect is causative or just correlative and to investigate the potential molecular mechanisms of this *trans*-effect.

Here we have shown that SylA treatment induces the expression of atypical RdDM components: *DRM1*, *AGO9* and *AGO3*. *DRM1* was previously shown to be mainly active during sexual reproduction (Jullien et al., 2012). Being preferentially involved in CHH methylation, the significant upregulation of DRM1 may at least in part explain the high abundance of CHH DMRs compared to the two other sequence contexts.

Interestingly, *AGO3* and *AGO9* were also characterized for their specificity to the reproduction phase of the *Arabidopsis* life cycle (Jullien et al., 2020; Olmedo-Monfil et al., 2010). So far, their role during bacterial infection remains to be investigated. The variation in genome-wide DNA methylation pattern upon proteasome inhibition by SylA could in part be linked to the up-regulation of these atypical RdDM components. Lastly, it is tempting to speculate that a DNA methylation reprogramming akin to the one happening during sexual reproduction might also happen during bacterial infection and be part of the plant defense mechanism.

## MATERIAL AND METHODS

### Plant materials and growth condition

Arabidopsis thaliana Wild-type Colombia-0 (*Col-0*) one week old seedlings were used throughout this study. Seeds were obtained from the NASC stock center (number N22681), stratified at 4°C for 2-3 days and germinated in sterile petri dish containing half Murashige and Skoog Basal Medium pH 5.7 (Ms ½, 0.8% micro agar and 0.215% MS in pH5.7 miliQ H2O). Seedlings were grown in the growth chamber under long-day conditions: 15 hours of light at 25°C and 60% of humidity and 10 hours of night at 21°C and 75% of humidity.

### Treatment with Syringolin A

Three to five one week old plantlets were transferred in a 24 wells plate with 500μL of liquid Ms ½. The plate was then return to the growth chamber and place under agitation. Seedlings were allowed to recover for 24h. After recovery, the seedlings were treated with Syringolin A (SylA) at a final concentration of 20μM during 24h.

Syringolin A stock solution (10mM in water) was stored at −20°C. After 24h hours, seedlings were collected and immediately frozen in liquid nitrogen and use for subsequent extraction.

### Sample preparation, qPCR and sequencing

Total RNA was extracted from seedlings using QIAzol Lysis Reagent (Qiagen). All samples were treated with DNase I (ThermoScientific) at 37°C for 30 minutes. DNAse I was subsequently inactivated by addition of EDTA and heat treatment (65°C for 10 minutes). First-strand cDNA were synthesized using 1yg of DNase treated total RNA and the Maxima First Strand cDNA Synthesis Kit (ThermoScientific), containing both oligo-dT and random hexamer primers. qPCR tests were performed with a QuantStudio 5 (ThermoScientific) using SYBR green (KAPA SYBR FAST qPCR Master Mix). qPCR mix were prepared according to the manufacturer’s protocol (KAPA Biosystems). An RNA equivalent of 25 ng of cDNA templates was distributed for each reaction. The qPCR program was as follow: 95 °C for 3 minutes followed by 45 cycles of 95 °C for 5 seconds and 60 °C for 30 seconds. ACTIN2 (AT3G18780) expression was used to normalize the transcript level in each sample. For each condition, RNA abundance of target genes was calculated from the average three independent biological replicates with three qPCR technical replicates. Real-time PCR primers used in this study are listed in Table S2.

For Genome wide bisulfite sequencing, seedlings genomic DNA was extracted using DNeasy Plant Mini Kit and was then processed and sequenced by Novogene (https://en.novogene.com/). Total sRNAs were trizol extracted and then processed into sequencing libraries and sequenced by Fasteris (http://www.fasteris.com, Switzerland). Total RNAs were extracted and DNAseI treated as previously mentioned. mRNA libraries were prepared and sequenced by Novogene (https://en.novogene.com/).

### Bioinformatics

#### mRNA profiling

Paired-end raw mRNA sequencing reads from two biological replicates per treatment regime were aligned to the *Arabidopsis thaliana* TAIR 10 genome assembly using HISAT2 (Kim et al, 2015), using default settings but setting maximal intron length at 10 kb. Aligned reads have been sorted using samtools sort command (Li et al., 2009). Aligned and sorted mRNA sequencing reads were mapped to genomic features using featureCounts (Liao et al., 2014). Multi-mapping reads were counted by assigning fractional counts to the respective features (“--fraction” option). Feature positional information was provided by a custom-made SAF file containing gene (and corresponding exon), transposable_element_gene, transposable_element, pseudogene, miRNA, ncRNA, snoRNA, tRNA, rRNA features retrieved from TAIR 10 gff annotion (TAIR10_GFF3_genes_transposons.gff obtained from www.arabidopsis.org). Differential expression has been analyzed using the R package edgeR (Robinson et al., 2010) using the exactTest function. Features with less than 15 reads across all samples were omitted. Differentially expressed features were defined by an absolute logFC > 2 and an adjusted *P*-value < 0.01. All plots have been generated using R-base functions. Gene ontology (GO) enrichment analysis has been performed using the R package topGO (Alexa et al., 2006) using GO term dataset “athaliana_eg_gene” retrieved form www.plants.ensembl.org. GO enrichment has been performed separately for up- and down-regulated DEGs.

#### small RNA profiling

Raw small RNA sequencing reads have been trimmed and subsequently filtered to remove reads mapping two rDNA loci (Chr2:1-10000 and Chr3:14194000-14204000), which have previously been repeatedly observed as a source of high number of sRNAs in *Arabidopsis thaliana* seedlings and, thus, compromise later analysis. Filtered sRNA reads were aligned to the Arabidopsis TAIR10 genome assembly using bowtie (Langmead et al., 2009) with the following options: -a --best --strata -m 10000. The aligned and sorted sRNA reads were split by size using a custom AWK script and were subsequently mapped to 100 bp genomic bins using HiCdat (Schmid et al., 2015). Only genomic bins, in which at least one of the samples had >= 10 reads were kept for further analysis. Both Ctrl and SylA had near identical effective library sizes (44810122 and 44827416 for sRNA between 17-30 nt), thus no further normalization has been performed. Genomic bins, which differed by more than 4-fold counts were considered differential sRNA bins for further analysis. To assign specific genomic features to differential sRNA bins, all non-ambiguous gff annotations have been extended by 1 kb up- and downstream of the start and end site and sRNA bins falling within these intervals were associated with the respective annotation unit. 5’ nucleotide identity has been assessed by a custom AWK script, taking into account all aligned sRNA reads of a given length within the respective 100 bp sRNA bins, whereas the sRNA read start had to lie within the bin. To associate differential sRNA bins with specific feature types, we employed bedtools intersect using custom-made bed files for different feature types, including promotor (defined as 1 kb upstream of TSSs of gene annotations), intergenic (excluding promotors), tRNA, miRNA, pseudogenes, TE (combination of transposable_element and transposable_element_gene, UTR (excluding promotor), intron, CDS (gene coordinates – introns). The initial coordinates for the features were retrieved from the publicly available TAIR10 gff file (see mRNA profiling).

#### Cytosine methylation analysis

Illumina sequencing from bisulfite-treated genomic DNA was aligned using bismark (Krueger and Andrews, 2011) with the following parameters: -q --score-min L,0,-0.2 --ignore-quals --no-mixed --no-discordant --dovetail --maxins 500. Sorted alignment files were analyzed for their cytosine methylation levels employing the R package methylKit (Akalin et al., 2012). Thereby, aligned reads were processed with the processBismarkAln() function setting the minimal coverage at 10 and minimal quality score at 20. Subsequently, methylation information per 100 bp genomic bin was generated using the tileMethyCounts() function. Differentially methylated regions (DMRs) were extracted using a q-value threshold of 0.05 and methylation difference cutoff of 10%. For further statistical analyses, methylated regions were defined with a cutoff of 40 % mC for CG context, 20 % for CHG context, and 10 % for CHH context. To associate a specific genomic feature (e.g., gene identifier) with a DMR, the DMR had to be positioned within a range of 1 kb 5’ of the annotation start to 1kb 3’ of the annotation end. To associate DMRs with specific feature types, we employed bedtools intersect using custom-made bed files for different feature types previously used to analyze feature type distribution in differential sRNA bins. Raw data are deposited on GEO under reference xxxx.

## ACKNOWLEDGMENTS

We would like to thank the following people for their help: Jasmine Sekulovski for support concerning plant growth, Gregory Schott and Olivier Voinnet for technical and financial support at the early stage of this study, Robert Dudler and Zsuzsanna Hasenkamp for providing Syringolin A.

## AUTHOR CONTRIBUTIONS

PEJ conceived the study. DMVB, LT and PEJ performed the experiments. SG performed the bioinformatic analysis. SG and PEJ analyzed the data. PEJ wrote the manuscript with the help of SG.

## Figure captions e-Xtras

**Fig. S1.**
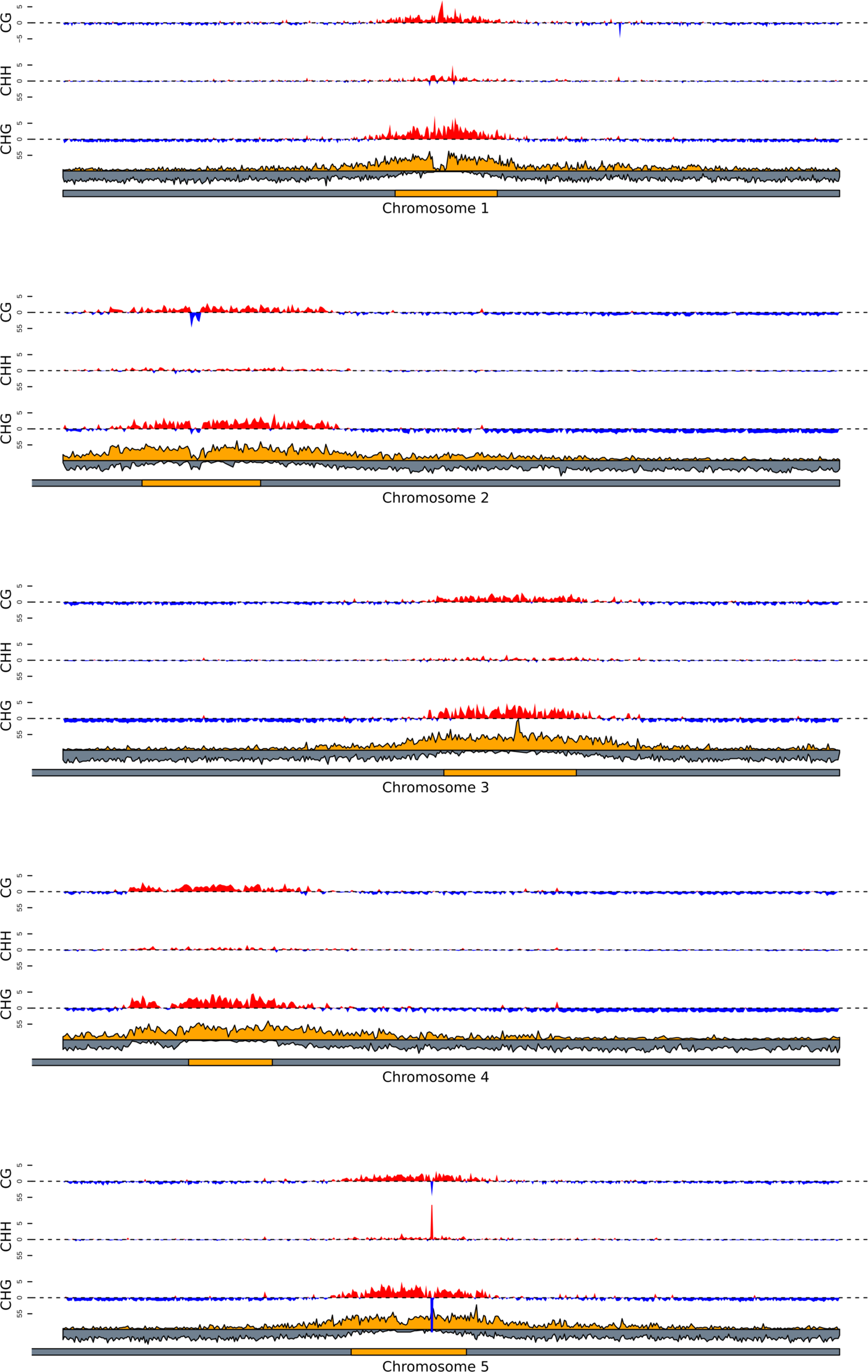
Percent methylation changes along chromosomes. Mean methylation change (in %) in 50 kb genomic bins, discerned by methylation context and *Arabidopsis* chromosomes 1-5. Red: hypermethylation, blue: hypomethylation. Gene (grey) and TE (yellow) densities in 50 kb are shown to highlight the occurrence of pericentromeres and chromosome arms, respectively.

**Fig. S2.**
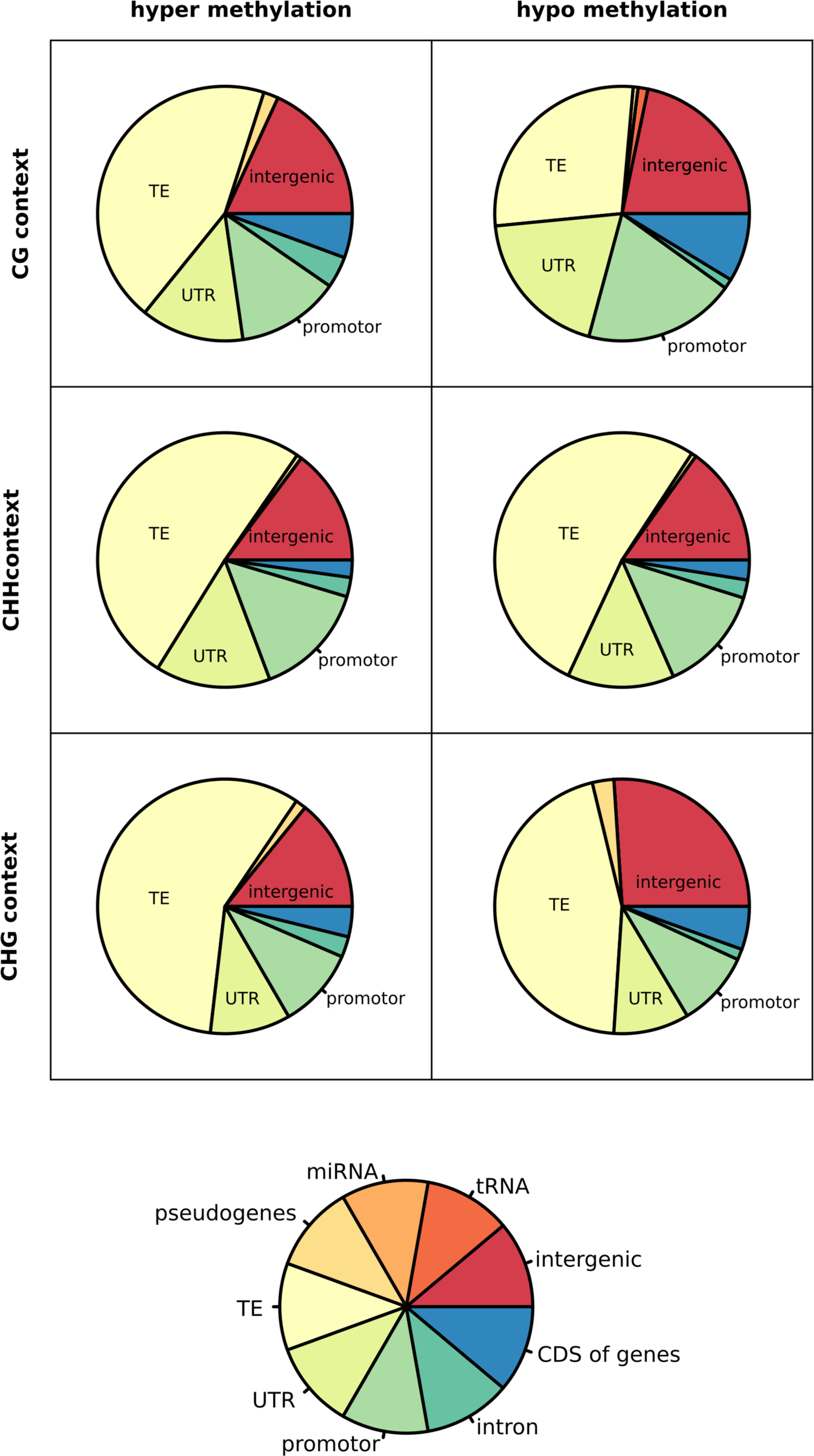
Genomic features associated to DMRs. Pie charts show the relative abundance of annotation feature types associated to a given type of DMR. Left: hypermethylated DMRs, right: hypomethylated DMRs, top: DMRs in CG context, middle: DMRs in CHH context, bottom: DMRs in CHG context.

**Fig. S3.**
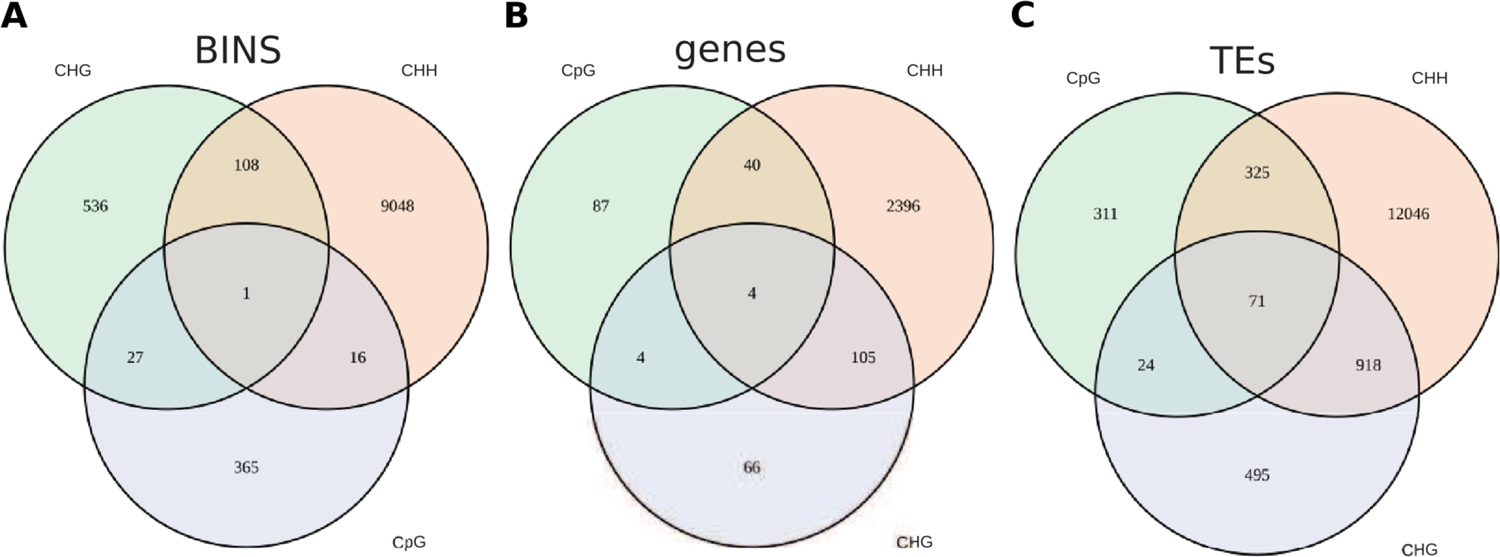
Overlaps between DMRs of different sequence contexts. (A-C) Venn diagrams showing overlap among the three sequence contexts (CHG, CHH, CG). A) Overlap among 100 bp genomic bins. B) Overlap among DMR-associated genes (associated with DMR, when DMR lies within – 1kb of the annotation start to +1 kb of the annotation end site) C) Overlap among DMR-associated TEs (associated with DMR, when DMR lies within – 1kb of the annotation start to +1 kb of the annotation end site).

**Fig. S4.**
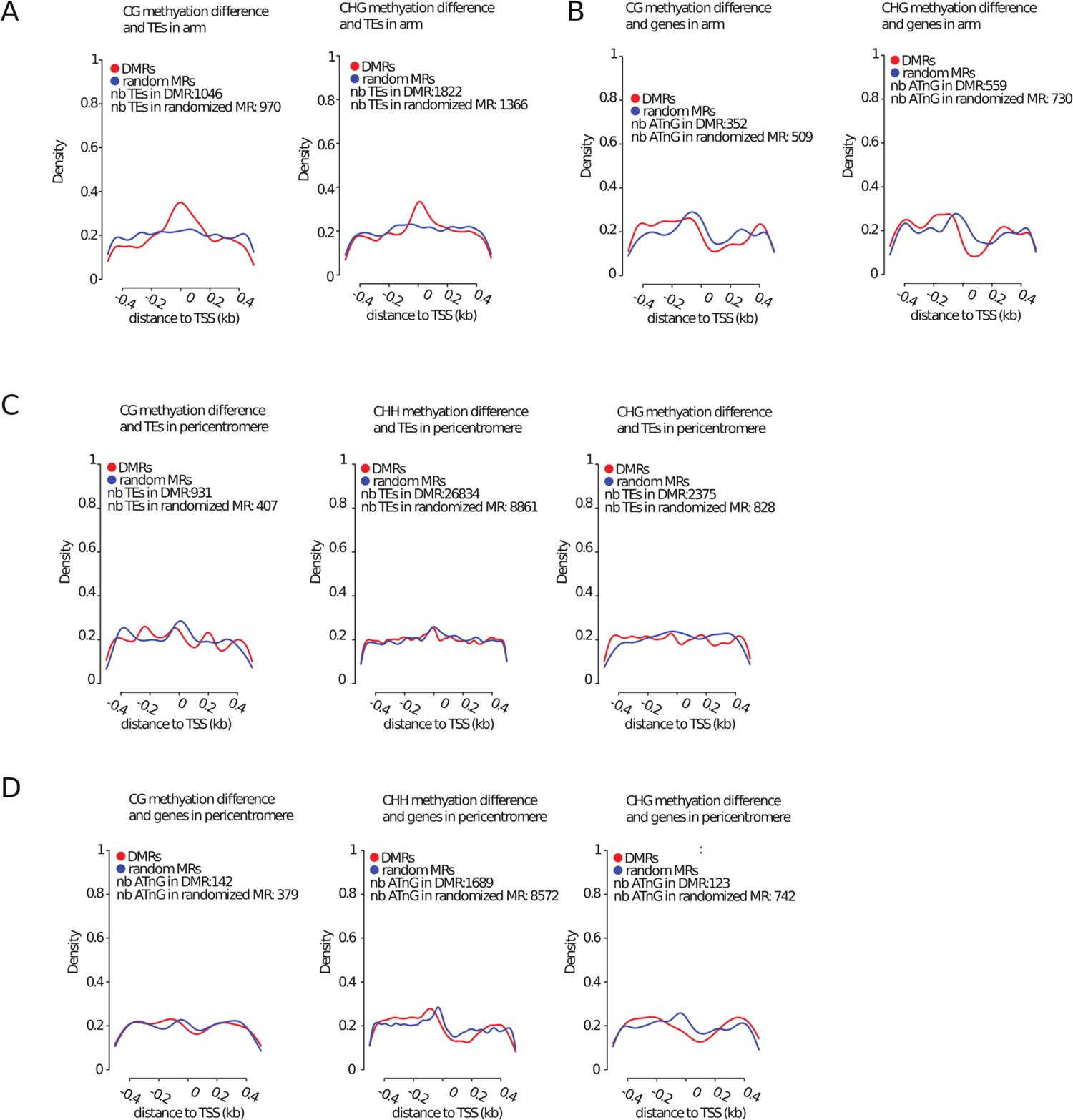
DMRs distance to transcription start site. (A-D) Metaplots representing the distribution of mean distance of DMRs to transcriptional start sites (TSSs). TSS closest to DMRs that lie within +/-5 kb of a TSS were selected. Red line: density of mean distance of DMRs to TSSs. Blue line: density of mean distance of randomly selected methylated regions (not being DMRs) that lie within +/- 5 kb of a TSS. A) TSS of TEs located on chromosome arms. B) TSS of genes located on chromosome arms. C) TSS of TEs located in pericentromeric region. D) TSS of genes located in pericentromeric region.

**Fig. S5.**
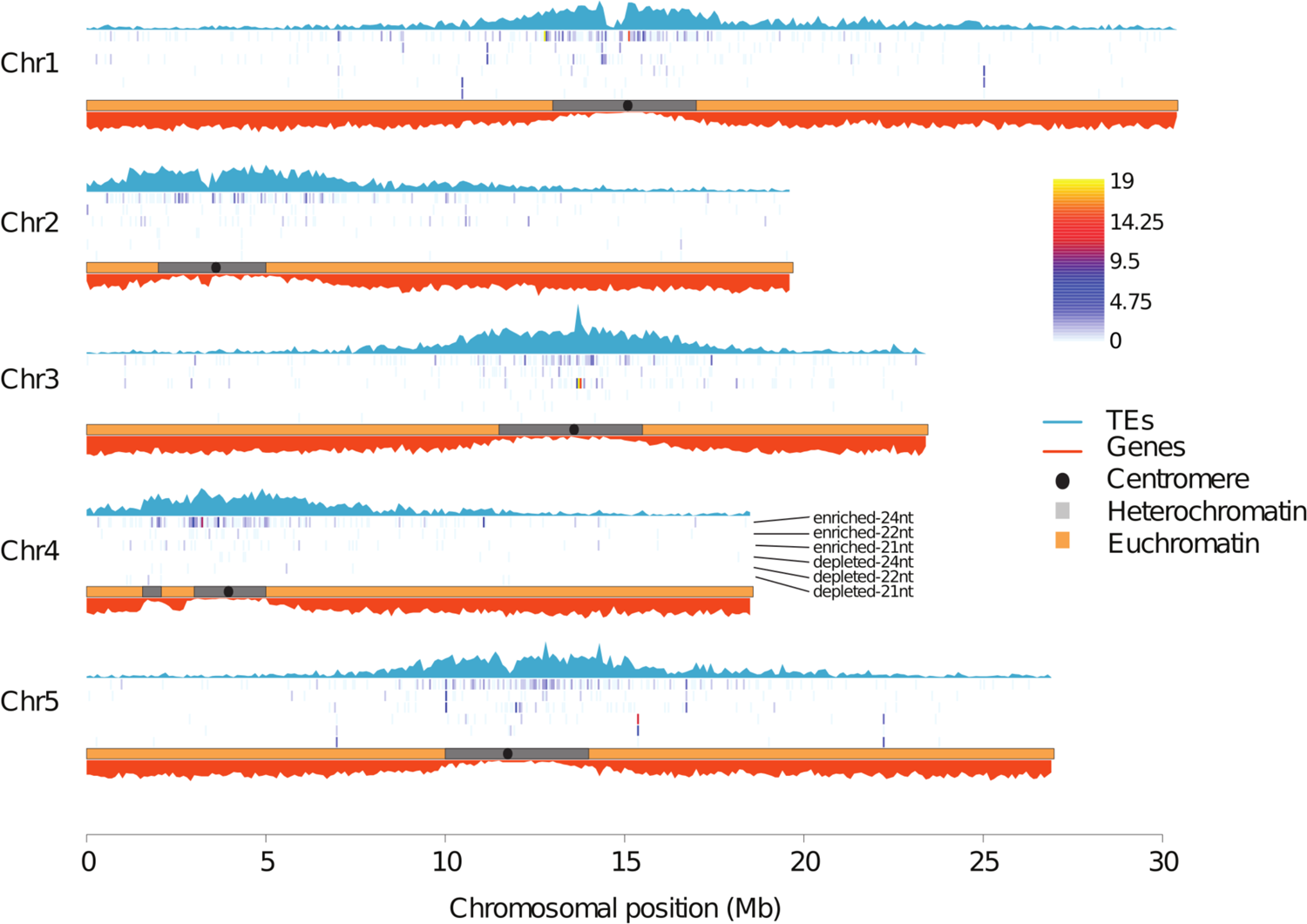
Genomic location of the differentially expressed sRNA bins. Chromosomal plot showing the number of differentially expressed sRNA bins along the five *Arabidopsis* chromosomes. The color code refers to the number of differentially expressed 100 bp sRNA bins within 50 kb genomic regions.

**Fig. S6.**
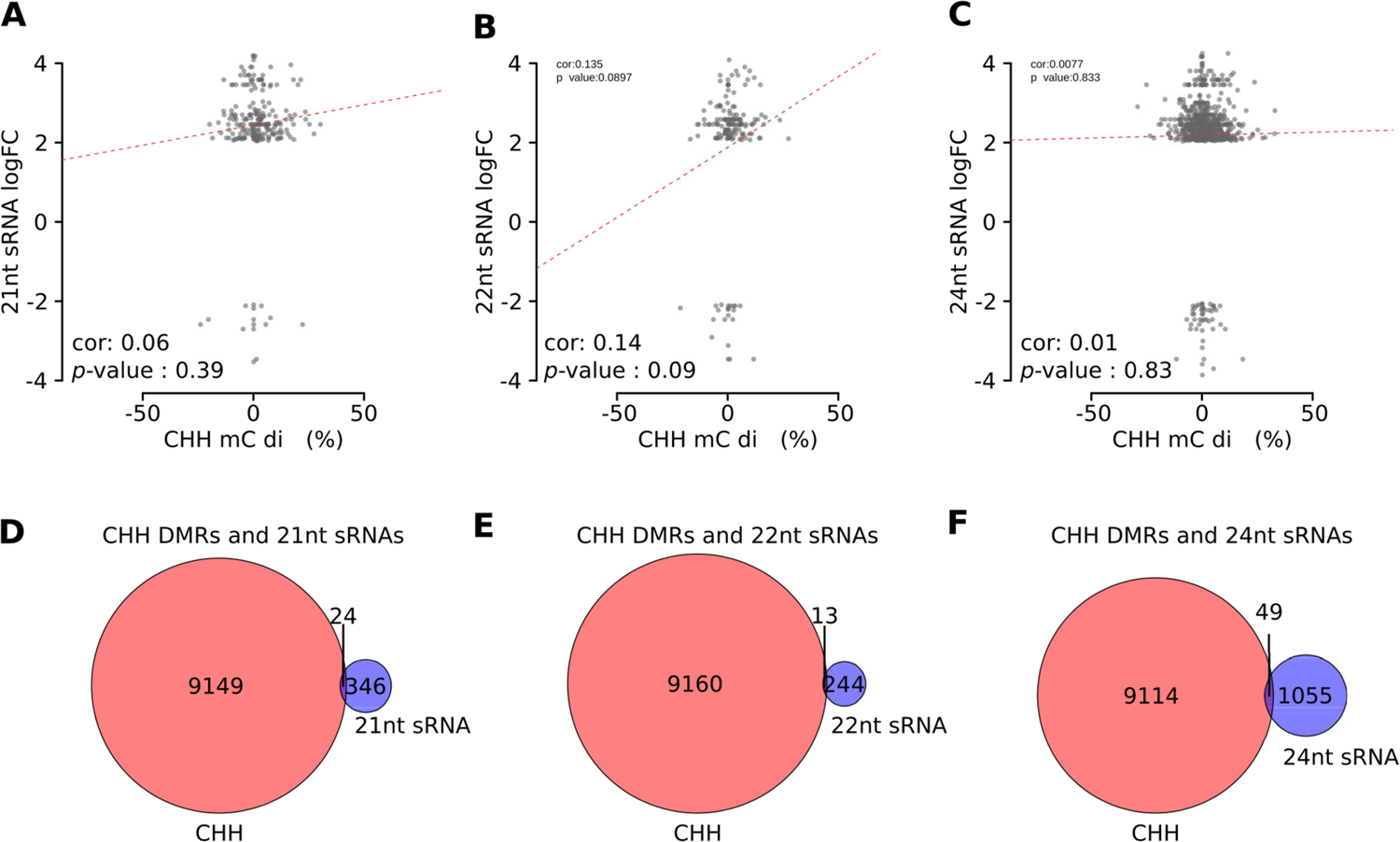
Relationship between differentially expressed sRNA bins and DNA methylation changes. (A-C) Scatterplots to visualize correlation analysis between differentially expressed sRNA bins and CHH methylation changes within these bins. A) 21nt sRNAs B) 22nt sRNAs C) 24nt sRNA. Test statistics: pearson correlation analyses. (D-F) Venn diagrams showing the overlap of 100bp CHH DMRs and differentially expressed 100bp sRNA bins. D) 21nt sRNA E) 22nt sRNA F) 24nt sRNA.

**Fig. S7.**
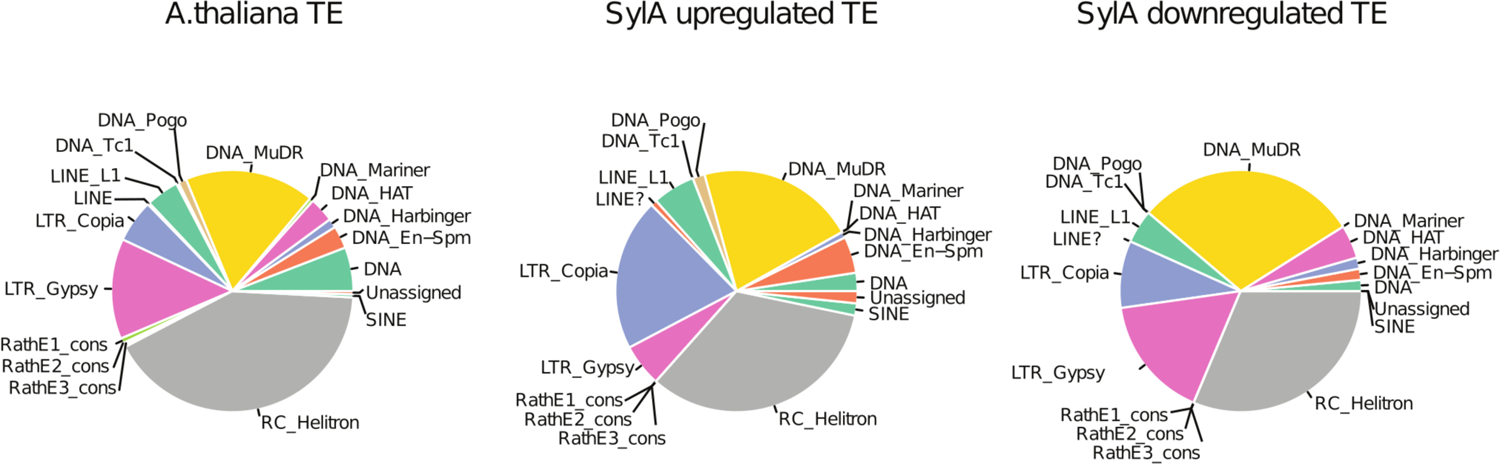
Differentially expressed TE families. Pie charts showing the relative abundance of TE families. A) Distribution of TE families in the *Arabidopsis thaliana* genome (TAIR10). B) Composition of TE families in significantly up-regulated TEs upon SylA treatment. C) Composition of TE families in significantly down-regulated TEs upon SylA treatment.

**Fig. S8.**
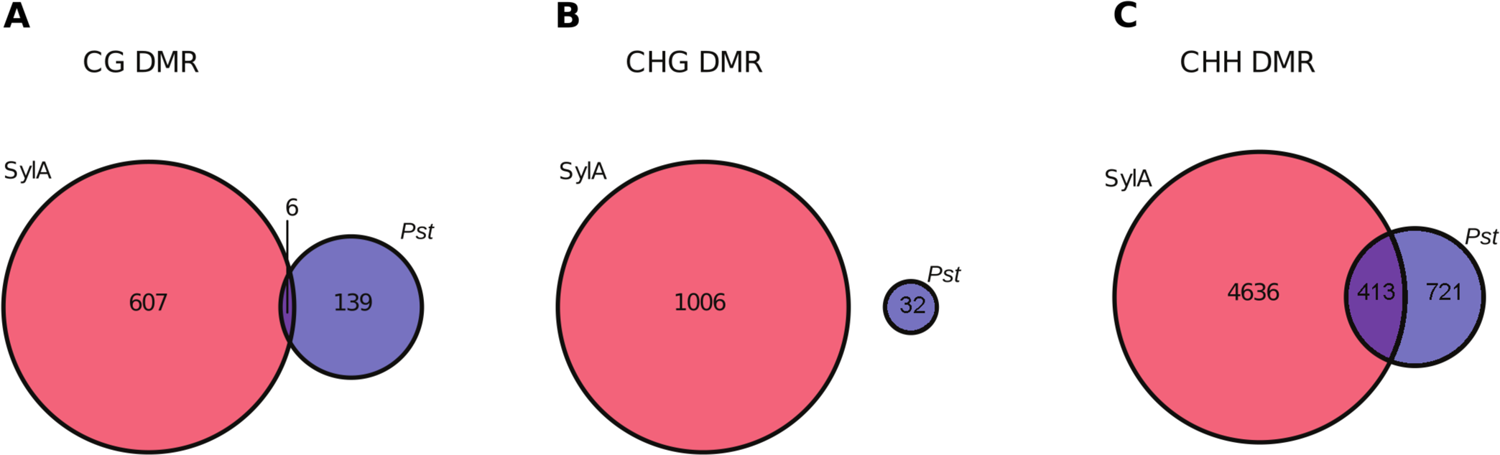
Overlap between SylA and *Pst* induced DMRs. Venn diagram representing overlapping DMR associated loci in SylA (this study) and *Pst* from the study by (Dowen *et al*. 2012) A) CG DMRs B) CHG DMRs C) CHH DMRs.

**Fig. S9.**
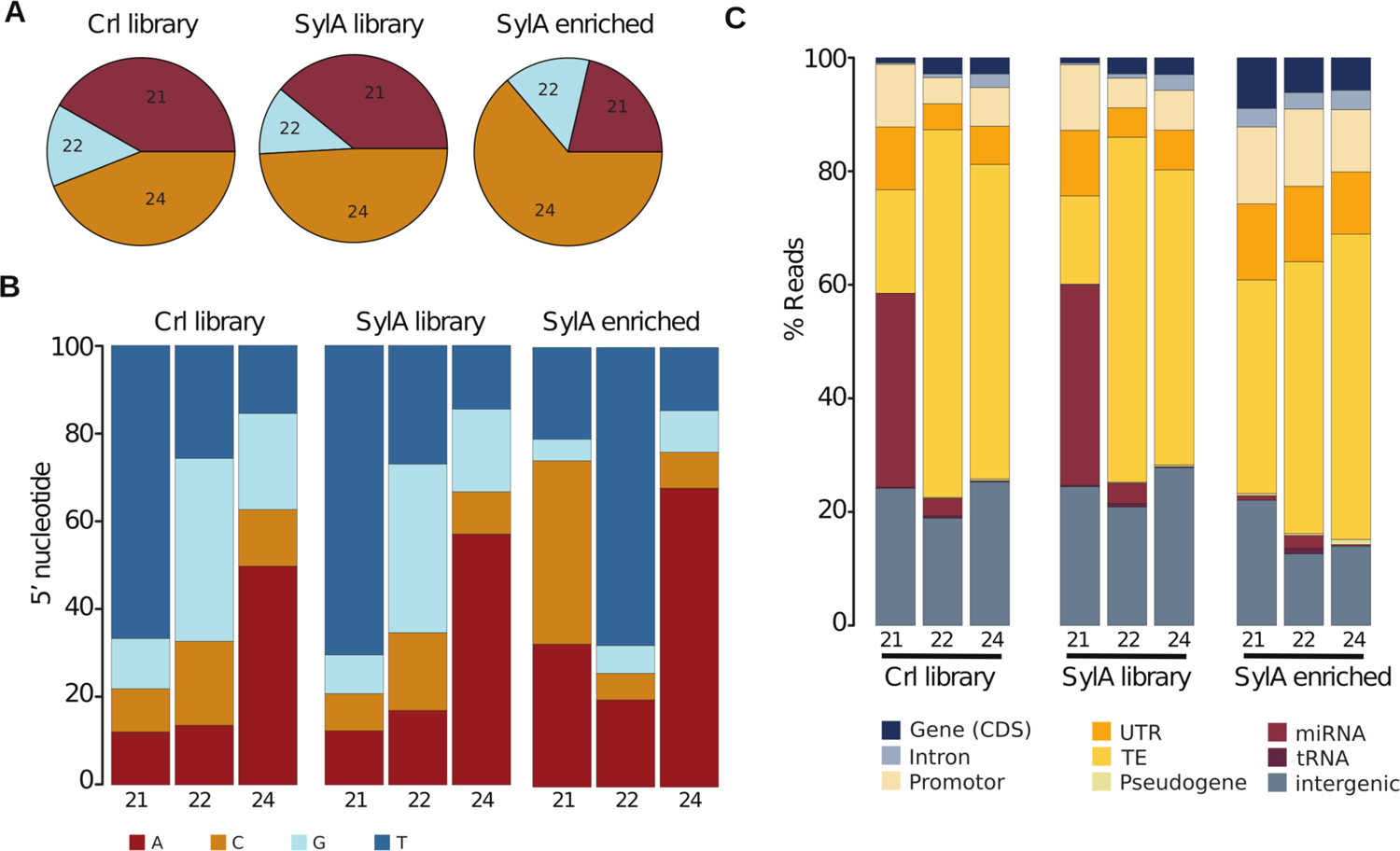
Characteristic of small RNA enriched population upon Syringolin A treatment. A) Pie charts representing sRNA size distribution (21 nt, 22 nt, 24 nt) in total mock (Ctrl) treated libraries and Syringolin A (SylA) treated libraries, and SylA enriched (logFC > 2) sRNA populations. B) Stacked bar charts showing the percentage of the 5’ nucleotide identity by sRNA sizes. C) Genomic features associated with sRNAs per size (21 nt, 22 nt, 24 nt). small RNA bins lying within 1 kb 5’ of the annotation start site and 1 kb 3’ of the annotation end site were selected.

